# Response of endolithic *Chroococcidiopsis* strains from the polyextreme Atacama Desert to light radiation

**DOI:** 10.1101/2020.09.01.278960

**Authors:** María Cristina Casero, Carmen Ascaso, Antonio Quesada, Hanna Mazur-Marzec, Jacek Wierzchos

**Affiliations:** Grupo de Ecología y Geomicrobiología, Dpto. Biogeoquímica y Ecología Microbiana, Museo Nacional de Ciencias Naturales, CSIC, Madrid, 28006, Spain; Departamento de Biología, Universidad Autónoma de Madrid, Madrid, 28014, Spain; Division of Marine Biotechnology, University of Gdansk, 81-378 Gdynia, Poland

**Author notes:** Correspondence to: María Cristina Casero.

## Abstract

The Atacama Desert is known to be the place on Earth with one of the highest solar radiation limiting the presence of life to endolithic microhabitats and soil microbial ecosystems. Endolithic microbial communities are supported by photosynthetic primary producers, mainly cyanobacteria, which can be injured by UVR. Nevertheless, cyanobacteria exposed to high solar radiation and its harmful effects have developed a series of defense mechanisms: avoidance, antioxidant systems or production of photoprotective compounds such as scytonemin among others. Scytonemin is a liposoluble pigment whose absorption maxima are located in UVA and UVC range and highly absorbing in the UVB range. In order to elucidate the protection capacity of endolithic cyanobacteria against harmful radiation, two cyanobacterial strains from *Chroococcidiopsis* genus were isolated from different endolithic microhabitats in the Atacama Desert: UAM813 strain, originally from the cryptoendolithic microhabitat of halite (NaCl), and UAM816 strain from chasmoendolithic microhabitat of calcite (CaCO_3_). Both were exposed to PAR and UVR+PAR conditions studying their short-term response, as oxidative stress and long-term response, as scytonemin production, metabolic activity and ultrastructural damage. The observed response of both strains reveals a high sensitivity to direct light exposure, even to PAR. The differences in their acclimation suggest specific adaptation strategies related to their original microhabitat, revealing their protective potential and the strain specific environmental pressure selection to inhabit different microhabitats and exposed to different light conditions.

**Importance:** Cyanobacteria are photosynthetic prokaryotes that inhabit most types of illuminated environments, even the endolithic microhabitats in cold and hot deserts. The environmental pressure caused by the extreme solar irradiation in the Atacama Desert involve that only those cyanobacterial strains able to cope with it can be found in these endolithic communities, usually dominated by members belonging to the extremotolerant *Chroococcidiopsis* genus. Here, a comprehensive analysis of multiple lines of defense against harmful sun radiation was conducted to diagnose the response of two *Chroococcidiopsis* strains isolated from different endolithic microhabitats and lithic substrates, and identify its relation with the original microenvironmental conditions of each strain. Our results contribute to a better understanding of the acclimation strategies developed by these cyanobacterial strains and its potential protective role for the whole endolithic microbial community.

## 1. Introduction

It is well known that ultraviolet radiation (UVR) at high dose might become lethal depending upon wavelengths, since some biomolecules can absorb harmful wavelengths (1). Absorption of those wavelengths may trigger an alteration in structure and activity of biomolecules, such as proteins, DNA and lipids, chronic depression of key physiological processes, and acute physiological stress leading to either reduction in growth and cell division, pigment bleaching, N_2_ metabolism, energy production, or photoinhibition of photosynthesis (2) (Fig.1). UVB radiation (280-320nm) has the greatest potential for cell damage since it has both direct effects on DNA and proteins (3). Likewise, UVA radiation (320–400 nm), produces indirect effects through the production of highly active oxidizing agents such as reactive oxygen species (ROS) (Fig.1).

**Figure 1.**
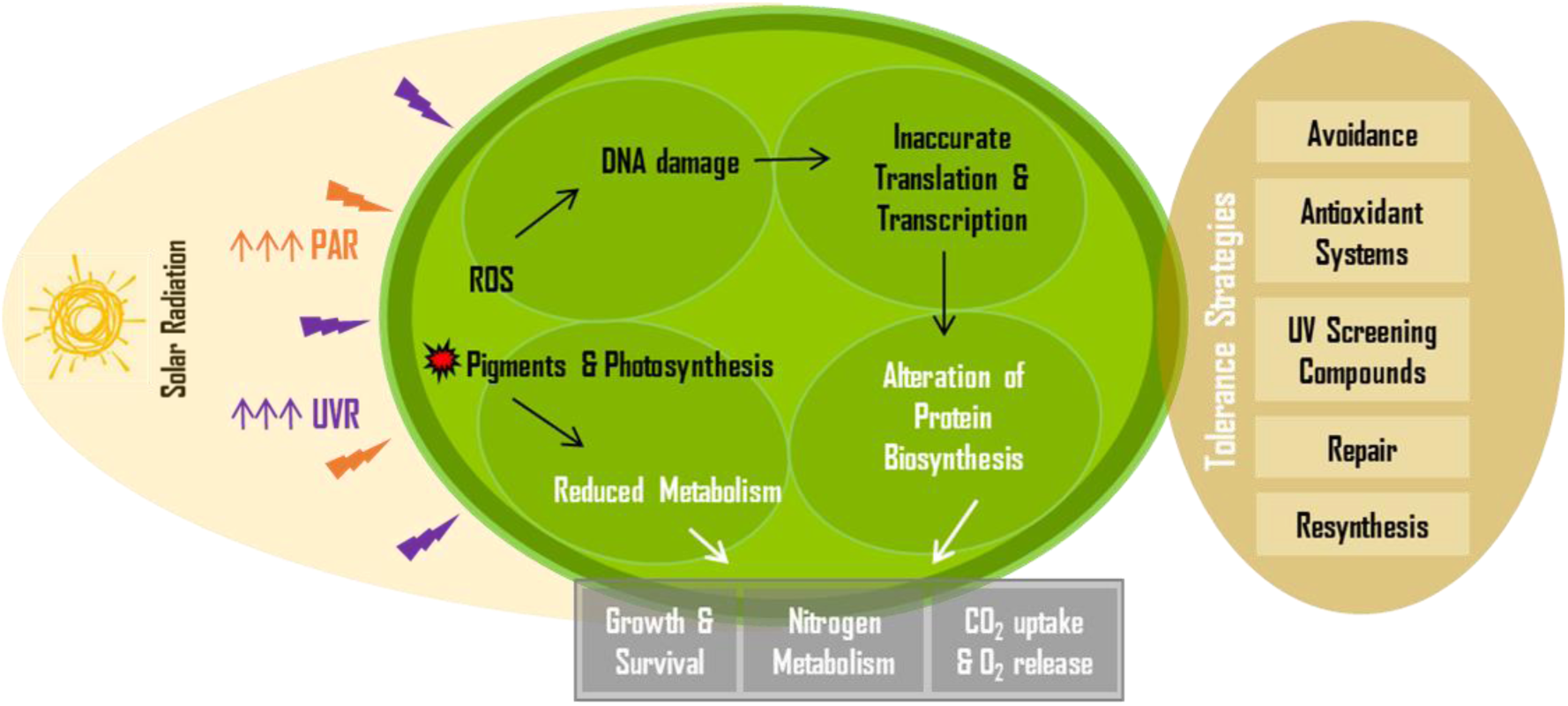
Summary of damaging effects, stress responses and tolerance strategies in cyanobacteria against long term exposure to extreme solar radiation. (modified from Rastogi et al. (14)).

The UVR-induced generation of ROS, such as hydroxyl radical (OH^-^), hydrogen peroxide (H_2_O_2_), singlet oxygen (^1^O_2_) and superoxide anion (O_2_^-^), has been reported in some cyanobacteria (4–6), yet very little is known about the subsequent effects of oxidative stress in these microorganisms. Oxidative stress produced by UVR can induce nucleotides modifications, translocations and DNA-DNA cross-links, an increase in the susceptibility to proteolysis and fragmentation of the peptide chain and oxidation of specific aminoacids (7). In fact, UVR affects genomic function and fidelity, as in the case of *Arthrospira platensis*, where an increase in thymine dimers frequency has been observed after a continued UVR exposure (8), in *Synechocystis* PCC 6308, whose DNA degradation due to UVR exposure has been reported (9), or in *Prochlorococcus marinus* PCC 9511, where UVR induced a delay in chromosomal replication (10). Furthermore, a differential lipid peroxidation in response to UVB exposure has been reported in *Nostoc muscorum, Plectonema boryanum* and *Aphanothece* (11), related to the oxidative degradation of polyunsaturated fatty acids in the cell membranes.

The photoinhibition process has been shown to occur when cyanobacteria are exposed to intense solar light, above the normal capacity of the photosynthetic electron flow (12) (Fig.1). Furthermore, different photosynthetic parameters, such as CO_2_ uptake, O_2_ evolution or ribulose-1,5 bisphosphate carboxylase (RuBisCo) activity are known to be inhibited when cyanobacteria are exposed to UVR (13). Likewise, UVB radiation is known to cause photobleaching of photosynthetic pigments such as chlorophyll *a* (14) and phycobiliproteins, generating a reduction on its content together with the disassembly of the phycobilisomal complex (15–17).

Since cyanobacteria originated in the Precambrian era, when the ozone shield was absent, UVR has presumably acted as an evolutionary pressure leading to the selection of UVR efficient protecting mechanisms (18), however, UVR tolerance varies between species. The mitigation strategies against the harmful effects of the exposure to UVR include avoidance, scavenging of ROS by antioxidant systems, the synthesis of UV-screening compounds, repairing systems for UV-induced DNA damage and protein resynthesis (14) (Fig.1).

Cyanobacteria rely on avoidance as a first line of defense against the potential damage caused by their exposure to UVR, preventing its harmful effects in different ways. Those inhabiting aquatic ecosystems can use migration to escape from high UVR to lower UVR intensities in the deeper par of water column (19), while some cyanobacterial species, especially terrestrial species, can move within the microbial mat structure by gliding to adapt their position to the required light regime, in order to minimize the harmful effects of intense solar light and UVR (17, 20), or colonize endolithic habitats (21).

As a second line of defense, cyanobacteria have developed complex antioxidant enzymatic or non-enzymatic systems to cope with UV-induced oxidative stress (22). Enzymatic antioxidants in cyanobacteria comprise superoxide dismutase (SOD), catalase (CAT), glutathione peroxidase (GPX) and the enzymes involved in the ascorbate-glutathione cycle. SOD protects different cellular proteins against oxidative stress and exists in four different metalloforms: Fe-SOD, Mn-SOD, Cu/Zn-SOD and Ni-SOD (23).

The third line of defense against UVR photodamage in cyanobacteria comprises the synthesis of UV-absorbing and/or UV-screening compounds (24, 25). Two main UV-absorbing/screening compounds known in cyanobacteria are: mycosporine-like amino acids (MAAs) and scytonemin. MAAs have an absorption spectrum from 310 to 362 nm and have found to be produced also by eukaryotic organisms such as fungi, microalgae, lichens and accumulated by invertebrate and vertebrate animals (26). They contribute to photostabilization and resist different physico-chemical stressors such as temperature, strong UVR and pH, becoming successful photoprotectants in diverse habitats (27). Several abiotic factors have been reported to affect the biosynthesis of MAAs in cyanobacteria (17), such as PAR and UVR (14), osmotic stress and desiccation (12, 28).

On the other hand, scytonemin is the most widespread sunscreen pigment, exclusively produced by cyanobacteria (12, 29). It is a yellow-brown lipid-soluble dimeric compound composed of indolic and phenolic subunits (30) and occurs in both oxidized (MW 544 Da) and reduced (MW 546 Da) forms. Its *in vivo* absorption maximum is at 370 nm and purified at 386 nm, in the UVA region. It shows high absorbance in the entire UVB region. This pigment is located in the EPS sheath of certain terrestrial cyanobacterial species is highly stable under different abiotic stresses being able to reduce about 90% of the UVA that reaches the cell (31). Due to high stability it can persist very long in terrestrial crusts or in dried mats (32) and performs its function without any further metabolic investment even under prolonged physiological inactivity.

The Atacama Desert has been pointed out as the place where some of the highest surface irradiance is likely to occur based on diverse features: its latitude (close to equator), its high altitude, relatively low ozone column values, prevalent cloudless conditions and low aerosol loading (33, 34). The World Health Organization uses the international standard measure, the UV index (UVI), to establish different risk levels of harm to humans so that regions where the UVI is greater than 11 would be positioned in the extreme risk of harm category. Following this criterion, the Atacama Desert has been described as the location on the Earth where the highest levels of surface UV irradiance have been measured with UVI reaching values up to 20 (35).

The extreme solar radiation is considered a limitation for life development, and thus not even epilithic (over the rock surfaces) microbial communities can be found in most of the hyper-arid region of the Atacama Desert due to the excessive exposure to the harmful effects of UVR (36). However, inhabiting endolithic microhabitats constitute an excellent first line of avoidance of the damaging effects of high radiation exposure for microorganisms in the Atacama Desert. The presence of few millimeters of lithic substrate over the endolithic microbial communities provides a certain barrier for UVR damage, as proposed by Cockell et al. (37), since only a 0.1-2.5% of the total incident solar radiation might reach the endolithic habitat depending on the substrate (38, 39).

Despite the UV-blocking effect provided by the lithic substrate used as an avoidance strategy, cyanobacteria from endolithic microbial communities in the Atacama Desert have been reported to exhibit the second line of defense, namely the production of carotenoids and possible antioxidant orange-carotenoid protein (40) as well as the third line of defense, the production of a UV-screening compound, such as scytonemin detected *in situ* by Raman spectroscopy (39–42).

*Chroococcidiopsis* species are considered extremotolerant organisms, occurring in a variety of terrestrial habitats. Members of the *Chroococcidiopsis* genus avoid high light intensities and UVR by living within soil, rock endolithic habitats and caves (21). Several strains from *Chroococcidiopsis* have been well characterized in order to identify their UVR tolerance by exposing *Chroococcidiopsis* cells to similar conditions as those occurring on Mars (43–45). Also the production of scytonemin, as the UVR defense mechanism of *Chroococcidiopsis*, was analyzed by Fleming and Castenholz (46) Dillon et al.(47) and Dillon and Castenholz (48).

The cryptoendolithic habitat in halite has previously been reported to harbor scytonemin produced by *Chroococcidiopsis* (42). Thus studying the response of the *Chroococcidiopsis* strain (UAM813) isolated from the translucent halite to direct solar simulated radiation gives an approach to its sensibility to this type of abiotic stress along with its capacity to protect the whole community. The second *Chroococcidiopsis* strain, isolated from the chasmoendolithic habitat in calcite (UAM816), was chosen since this type of endolithic habitat is more exposed to direct solar radiation. Hence this works aims to diagnose the stress response to UVR and PAR of *Chroococcidiopsis* strains isolated from two different endolithic microhabitats from the region with one of the highest solar radiation on Earth, in order to identify its relation with features of their original microenvironmental conditions.

## 2. Materials and Methods

### 2.1. Culture organisms and conditions

Two strains of cyanobacteria isolated from endolithic habitats of the Atacama Desert were used in this study: *Chroococcidiopsis* UAM813, from the cryptoendolithic microhabitat in halite from Yungay area (24°05’09’’ S, 069°55’17’’ W) and *Chroococcidiopsis* UAM816, from the chasmoendolithic microhabitat in calcite from Valle de la Luna (22°54’39’’ S, 06814’49’’ W). Both strains are preserved at the Universidad Autónoma de Madrid, Madrid, Spain. *Chroococcidiopsis* UAM813 and UAM816 were grown as batch cultures in BG11 medium (49) at 28°C under continuous 12 W m^-2^ PAR (∼ 60 µmol photons m^-2^ s^-1^) generated by cool white fluorescent lamps.

All described experiments were performed in triplicates following the Fleming and Castenholz (46) indications. The experimental design remained as follows: cultures were gently homogenized by orbital shaking. Three milliliter aliquots of the homogenized cultures were then filtered onto 25 mm diameter, 0.2 µm pore size Cyclopore Track-Etch Membranes (Whatman), producing a thin layer of cells on the filter. The filtered cells were immediately transferred to 1% agar plates made with BG11 medium. The agar plates with the filters and filtered cells were then kept under the following conditions: continuous 40 W m^-2^ PAR at 25°C, for 2 days. At the end of the 48 h period, a UVA lamp (F20T10/BLB lamp (315-400 nm)) was turned on exposing half of the entire set of filtered cells to continuous 2 W m^-2^ UVA radiation in addition to PAR.

It is assumed that the experimental conditions described above are definitely not exactly the same as microenvironmental conditions within the cryptoendolithic (halite) or chasmoendolithic (calcite) habitats. However, this experimental design is approach to UVA+PAR irradiance within the endolithic habitat in the Atacama Desert.

PAR (400-700 nm) and UVR (215-400 nm) measurements were made using an ULM-500 universal light meter (Heinz Walz GmbH, Effeltrich, Germany) and Apogee UV Radiation MU-200 meter, respectively. All readings refer to values measured on the surface of the cultures.

### 2.2. *In vivo* detection of oxidative stress

The spectrophotometric detection of the production of ROS after defined time intervals (24, 48 and 72 hours) of exposure to simulated solar radiation was performed by using 2’,7’-Dichlorodihydrofluorescein diacetate (DCFH-DA) (Sigma Aldrich-Merck KGaA, Darmstadtm Germany) solubilized in ethanol. Filtered cells were resuspended in 1 mL phosphate buffer (PBS) where a 5 µM (final concentration) of DCFH-DA was added. Samples were then incubated in a shaker at room temperature in the dark for 1 h. DCFH is nonfluorescent but switched to highly fluorescent DCF when oxidized by intracellular ROS or other peroxides having an excitation wavelength of 485 nm and an emission band between 500 and 600 nm. After 1 h incubation, samples were subjected to fluorescence spectrophotometric analysis. The fluorescence of the samples was measured by a spectrofluorophotometer with an excitation wavelength of 485 nm and an emission band between 500 and 600 nm. The fluorescence intensity was corrected against the blank control experiments without cells and then normalized to dry weight. Its comparison with control samples was used to determine the oxidative stress. All fluorescence measurements were performed in triplicates. Every measurement was normalized to dry weight using a XP6 microbalance (Mettler Toledo, Columbus, OH, USA).

CellROX Green reagent (Invitrogen) was used to detect ROS in both *Chroococcidiopsis* by fluorescence microscopy following the optimized method described by Cornejo-Corona et al. (50). Briefly, 2 µL of 5 mM CellROX Green was added to 100 µL of cyanobacterial culture followed by incubation at room temperature and shaking at 120 rpm for 30 min in the dark. The cells were then washed twice for 5 min, each time at room temperature with 1× PBS, 0.1% Triton X-100, and fluorescence was observed using a Zeiss AxioImager M.2 fluorescence microscope (Carl Zeiss, Jena, Germany) and a Apochrome x60, n=1.4 Zeiss oil-immersion objective. Multichannel Image Acquisition (MIA) system was used with a combination of the following filter sets: filter set for eGFP (Zeiss Filter Set 38; Ex/Em: 450-490/500-550 nm) for CellROX green fluorescence and weak EPS autofluorescence signal, and Rhodamine (Zeiss Filter Set 20; Ex/Em: 540-552/567-647 nm) for red chlorophyll *a* and phycobiliproteins autofluorescence signal. At least one hundred cells were evaluated for each experimental time and treatment. The samples were observed under bright field light to locate aggregates for evaluation and then the microscope was switched to fluorescence to identify the number of fluorescent cells and their signals.

Although cultures were not axenic, heterotrophic biomass never exceeded 1-2% of the total biovolume based on cell counts according to Schallenberg et al. (51).

### 2.3. Scytonemin induction experiment

Filtered *Chroococcidiopsis* UAM813 and UAM816 cells were exposed to PAR or UVR+PAR light in two separate sets for 3, 6, 9, 12 and 15 days. After the respective experimental exposure time cells were scraped out of the filters and suspended in the BG11 medium and then gently homogenized by pumping them multiple times with a 1000 µl Pipetman (Gilson, Middleton, WI, USA). For the determination of scytonemin content, cells were suspended in 1:1 (v/v) methanol: ethyl acetate (M-EA) by overnight incubation at 4°C in darkness (29). After centrifugation (10,000x *g* for 5 min) samples were filtered through 0.2 µm pore-sized sterilized syringe-driven filter (Symta, Madrid, Spain) before being subjected to HPLC analyses.

Partially purified scytonemin was analyzed using a HPLC system (Agilent Technologies 1200 Series, Photodiode Array). The 20 μL of M-EA extract were injected into the HPLC column Phenomenex Peptide 100 Å, 3.6 μ × 4.60 mm; XB C18. Elution was at a flow rate of 0.5 mL min^-1^ using the mobile phase composed of solvent A (5% acetonitrile in milliQ water + 0.1% formic acid) and solvent B (100% acetonitrile + 0.1% formic acid). The 30 min gradient elution program was set with 0–15 min linear increase from 15 % solvent A to 80 % solvent B, and 15–30 min at 100 % solvent B. The detection wavelength was at 384 nm. The PDA scan wavelength ranged from 200 to 700 nm. Oxidized and reduced scytonemin were identified by their characteristic absorption maxima.

At the same time, the absorbance of each extract was measured at 384 (scytonemin maximum), 490 (pooled carotenoids) and 663 nm (Chl *a*). These absorbance values were partially corrected for residual scatter by subtracting the absorbance at 750 nm. Absorbance measurements were made on a Flame Spectrometer (Ocean Optics, Florida, US.).

Every measurement was normalized to dry weight using a XP6 microbalance (Mettler Toledo, Columbus, OH, USA). Although cultures were not axenic, heterotrophic biomass never exceeded 1-2% of the total biovolume based on cell counts according to Schallenberg et al. (51).

### 2.4. Metabolic Activity experiment and UVR effect on *Chroococcidiopsis* cellular ultrastructure

The metabolic activity of *Chroococcidiopsis* cells was evaluated at 6 experimental times (0, 3, 6, 9, 12, 15 days) for both experimental conditions, PAR and UVR+PAR. For this purpose, the cell-permeable 5-Cyano-2,3-Ditolyl Tetrazolium Chloride (CTC) redox dye was used. This dye is reduced from a soluble colorless form into its corresponding fluorescent insoluble formazan (CTF) that accumulate intracellularly. The formazan crystals are viewed as intracellular opaque dark-red deposits under bright field illumination, or as yellow-orange fluorescent spots (excitation and emission maxima at 488 and 630 nm) when using fluorescence microscopy.

Procedures described by Tashyreva et al. (52) were followed for CTC staining, increasing incubation times from 2 to 5 hours. A Zeiss fluorescence microscope (AxioImager M2, Carl Zeiss, Germany) was used with Apochrome oil immersion objective x64, n=1.4. The optical system for CTF fluorescence observations included HE Rhodamine filter set (Ex/Em: 426-446 / 545-645 nm).

### 2.5. Light microscopy

Light microscopy in differential interference contrast (DIC) mode was performed on cell aggregates of both *Chroococcidiopsis* strains at each experimental time for the scytonemin induction experiment. The samples were examined using a microscope (AxioImager M2, Carl Zeiss, Germany) equipped with Apochrome x64, n=1.4 oil immersion objective.

### 2.6. Transmission Electron Microscopy (TEM)

Cyanobacterial cells from UAM813 and UAM816 *Chroococcidiopsis* strains were centrifuged at 3,000x *g* and resuspended in 3% glutaraldehyde in 0.1M cacodylate buffer and incubated at 4°C for 3 hours. The cells were then washed three times in cacodylate buffer, postfixed in 1% osmium tetroxide for 5 hours, before being dehydrated in a graded series of ethanol and embedded in LR White resin (53). Ultrathin sections were stained with lead citrate and observed with a JEOL JEM-2100 electron microscope (Tokio, Japan) equipped with Gatan Orius CCD camera (Pleasatan, CA, USA) at 200kV acceleration potential.

### 2.7. Statistical analysis

All results are presented as mean values of three replicates. Data from scytonemin induced production and oxidative stress were analyzed by one-way analysis of variance. Once a significant difference was detected post-hoc multiple comparisons were made by using the Tukey test. The level of significance was set at 0.05, 0.01 and 0.001 for all tests.

## 3. Results

### 3.1. Oxidative stress in *Chroococcidiopsis*

#### Semi quantitative analysis of intracellular ROS by DCF fluorescence

Oxidative stress on both *Chroococcidiopsis* strains was examined in vivo by DCF fluorescence normalized to dry weight at 4 experimental times for a 72-hour period. The strains were exposed to two different light conditions: to 40 Wm^-2^ PAR, or under that PAR together with 2 Wm^-2^ UVR (UVR+PAR). ROS accumulation, represented by DCF fluorescence, in the UAM813 strain increased after 24 hours of exposure, increasing sequentially and reaching maximum fluorescence after 72 hours of exposure (Fig. 2). Light treatment revealed a significant effect on ROS accumulation (R^2^, 0.979) and punctual significant differences were observed at all three experimental times. Specifically, PAR conditions revealed a higher accumulation of ROS after 24 and 72 hours of exposure, while UVR+PAR conditions involved significantly greater ROS accumulation after 48 hours of exposure (Fig. 2).

**Figure 2.**
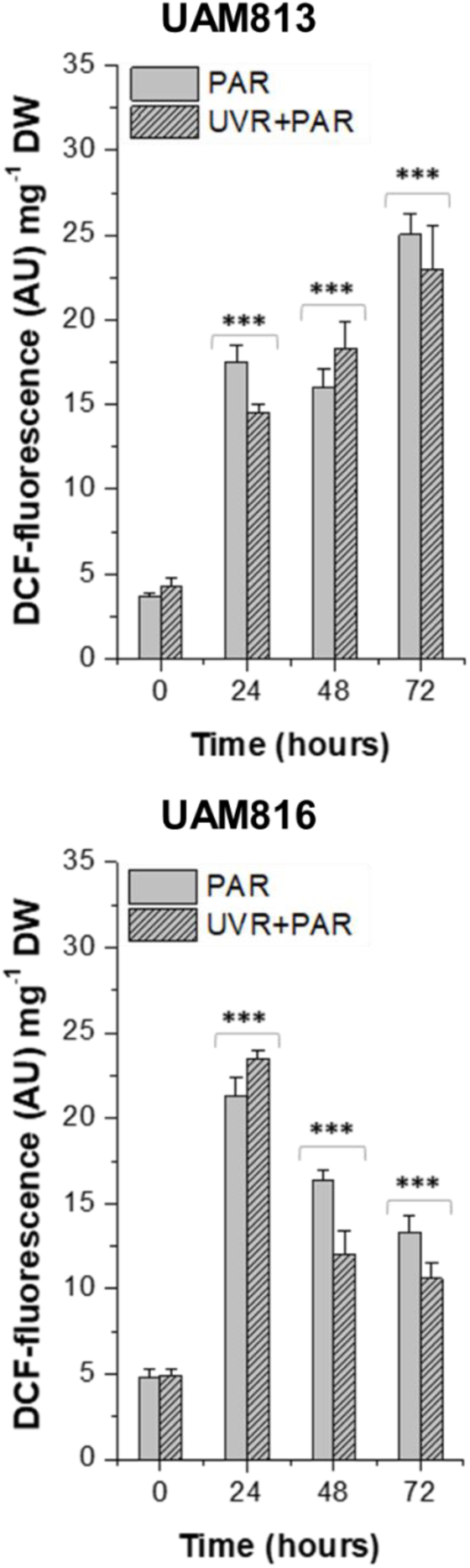
DCF fluorescence measurements in *Chroococcidiopsis* strains UAM813 (upper graph) and UAM816(lower graph) after irradiation with PAR (plain bars)or UVR+PAR (scratched bars) for 72h normalized to dry weight. Significant differences between light conditions at marked by *** (0.001).

Oxidative stress response in the UAM816 strain shared an increase after 24 hours of exposure for both light treatments where maximum DCF fluorescence values were registered. A significant decrease in DCF fluorescence was observed after 48 and 72 hours. This strain showed a significantly lower accumulation of ROS after 24 hours of PAR in comparison to UVR+PAR conditions. By contrast, significant differences observed at 48 and 72 hours of exposure occurred due to a higher ROS accumulation after PAR exposure (Fig. 2).

#### Reactive oxygen species formation

The visualization of ROS produced by *Chroococcidiopsis* under PAR and UVR stress was performed based on the dye CellROX Green at time 0 and after 24 hours of exposure to UVR+PAR on both strains (Fig. 3).

**Figure 3.**
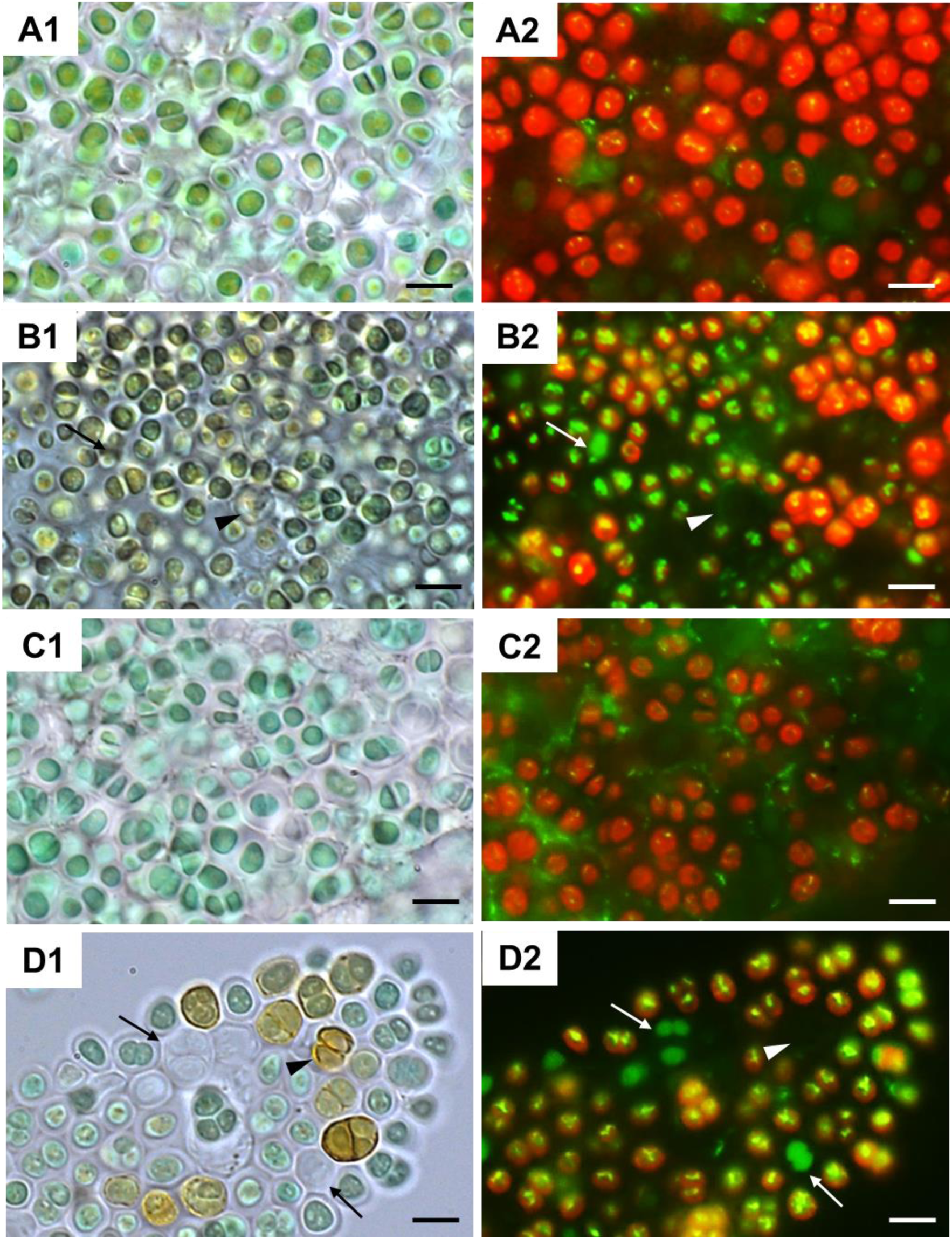
CellRox staining for intracellular detection of ROS in *Chroococcidiopsis* strains UAM813 (Series A and B) and UAM816. Series C and D). Bright field and fluorescence microscopy images of Series A and C correspond to exposure time t=0, and from Series B and D after 24 hours of exposure to UVR+PAR. Red signal correspond to cyanobacteria chlorophyll and phycobiliproteins autofluorescence, higher in non-exposed cells (A2, C2) than in UVR+PAR exposed cells (B2, D2). Bright yellow/green dots in fluorescence images are due to CellRox fluorescence, the oxidative stress indicator, higher in UVR+PAR exposed cells (B2, D2) than in non-exposed cells (A2, C2). On images of UVR+PAR treated cells, arrows point to cells revealing apparently structural integrity (B1 and D1) with green autofluorescence signal (B2, D2). In images of UVR+PAR treated cells, arrow heads point at cells revealing apparently structural integrity and brown color (B1, D1), suggesting an increase in scytonemin content, and no autofluorescence signal (B2, D2). Scale bars = 8 μm.

Bright field microscopy images on Figs. 3 at time 0 (Figs. 3A1 and C1) and 24 hours of exposition to UVR+PAR (Figs. B1 and D1) already exhibited differences in the color of the cells. The UAM813 strain and UAM816 strain were light green and blue-green respectively at time 0 turning to brownish green and yellow-brown color after 24 hours of exposure to UVR+PAR light conditions.

Fluorescence microscopy images on Figs. 3 shown intense red autofluorescence of chlorophyll and phycobiliproteins of *Chroococcidiopsis* cells for both UAM813 (Fig. 3A2) and UAM816 (Fig. 3C2) strains at time 0. The weak green autofluorescence of EPS signal could be observed outside the cells for both strains at time 0. After 24 hours of UVR+PAR exposure the oxidation of CellROX fluorochrome by ROS and its binding to DNA lead to the formation of bright spot-like green-yellow fluorescence inside the UAM813 (Fig. 3B2) and UAM816 (Fig. 3D2) respectively.

Moreover, many of both *Chroococcidiopsis* strain cells after their irradiation for 24 hours clearly lost the red autofluorescence emitting only green autofluorescence signal (white arrows in Figs. 3B2 and D2). Some of the cells even did not reveal any fluorescence signal (white arrow heads on Figs. 3B2 and D2). These cells appear as cell-shaped brown covers on bright field images (black arrow heads on Figs 3B1 and 3D1)

### 3.2. Scytonemin induction in *Chroococcidiopsis*

The total scytonemin content in the *Chroococcidiopsis* strains UAM813 and UAM816 was evaluated using two different methodologies such as, HPLC quantification and trichromatic equation based on UV-VIS absorption spectra. No statistical differences in scytonemin values were found between both quantification methods.

The scytonemin content in UAM813 strain (Fig. 4) increases with time during the first 9 days of experiment in both experimental conditions. When this strain was exposed only to PAR light, the maximum content of 16.4 μg scytonemin mg^-1^ DW was detected after 9 days of exposure. When the strain was exposed to UVR and PAR light, maximum scytonemin content of 20.8 μg mg^-1^ DW, was detected also after 9 days of exposure. Significant differences between both experimental conditions were observed after only 6 days of exposure, where the scytonemin content under UVR+PAR conditions was 14.6 μg mg^-1^ DW, while it remained as low as 1.9 μg mg^-1^ DW under PAR only. Significant differences in scytonemin content between both light treatments were also observed during the last 6 days of experiment. The scytonemin content decreased drastically after 12 days of experiment under PAR light down to the initial values at time 0. However, the tendency was significantly different when UVR+PAR light was used, finding non-significant differences in scytonemin content between the last experimental time and the value detected after 9 days.

**Figure 4.**
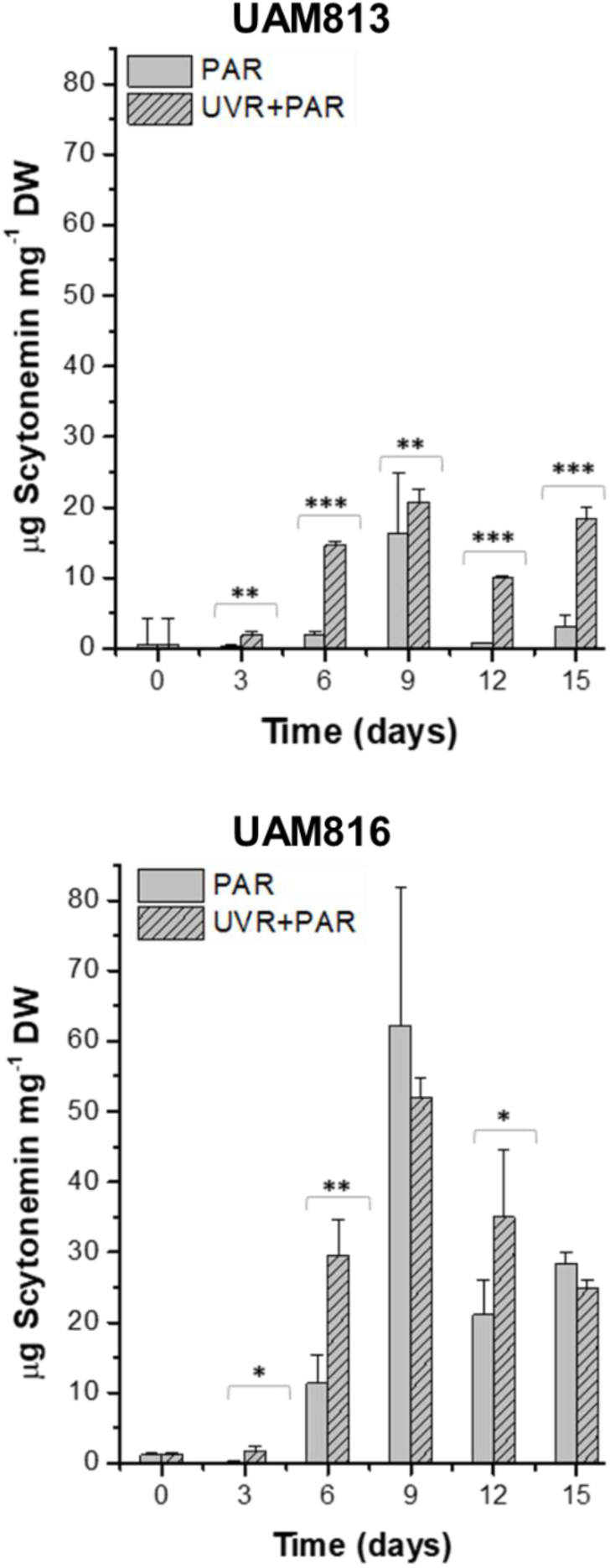
Total scytonemin content on *Chroococcidiopsis* strains UAM813(upper graph) and UAM813(lower graph) after irradiation with PAR (plain bars)or UVR+PARR(scratched bars) for 15 days normalized to dry weight quantified by HPLC. Significant differences between light conditions are marked by *** (0.001); ** (0.01); * (0.05).

The maximum scytonemin content in the UAM816 strain (Fig. 4) was reached after 9 days of exposure to both experimental conditions, with no significant differences between them. This maximum content was 62.3 μg scytonemin mg^-1^ DW for PAR light conditions and 52 μg scytonemin mg^-1^ DW for UVR+PAR light conditions. Relative scytonemin content under both light conditions decreased during the last 6 days of exposure. Significant differences were observed between the experimental conditions at three different times, after 3, 6 and 12 days of exposure. The greater difference between both treatments occurred after 6 days of exposure, exhibiting an almost three times higher scytonemin content after UVR+PAR treatment (29.6 μg mg^-1^ DW) in comparison with PAR treatment (11.4 μg mg^-1^ DW).

#### Scytonemin characterization

A further analysis was performed to partially characterize scytonemin. The HPLC analysis of the scytonemin showed two prominent peaks in chromatograms of both *Chroococcidiopsis* strains (Figs. S1, A-B) at retention time 16.57 min (a) and 17.89 min (b) with a UV absorption maximum at 385 nm identified as reduced scytonemin (a) and oxidized scytonemin (b) according to Rastogi and Incharoensakdi (29). The obtained chromatogram revealed the presence of both reduced and oxidized scytonemin in the M-EA extracts of both *Chroococcidiopsis* strains. However, the proportion of each type of scytonemin accumulated by both *Chroococcidiopsis* strains cannot be revealed due to the presence of O_2_ in atmosphere during the extraction procedure and possible partial oxidation of reduced scytonemin.

### 3.3. Metabolic activity of *Chroococcidiopsis*

#### Chroococcidiopsis *cells metabolic activity analysis*

The metabolic activity of the *Chroococcidiopsis* cells was evaluated after 15 days of exposure to two different light conditions, PAR and UVR+PAR.

Three categories were established for the vital status of the *Chroococcidiopsis* cells depending on their fluorescence emission after the CTC staining. Those with green autofluorescence (GF+) were defined as not viable cells according to Roldan et al. (54). The cells exhibiting only red chlorophyll and phycobiliproteins autofluorescence (PCHL+) were defined as damaged (not metabolically active); while cells presenting both chlorophyll and phycobiliproteins red autofluorescence and bright orange spots (CTF crystals) (PCHL+ / CTC+) were defined as metabolically active (Fig. 5).

**Figure 5.**
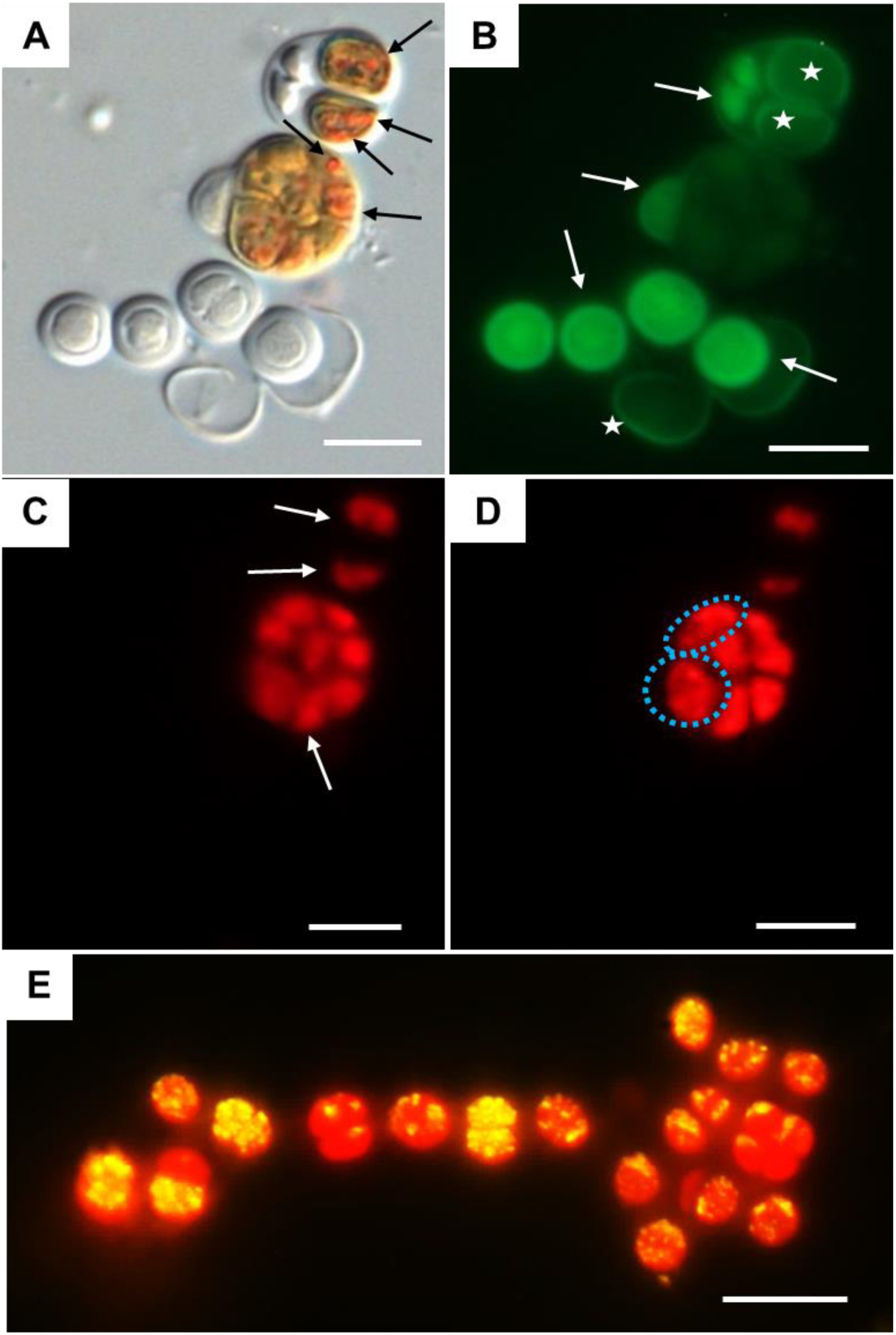
DIC and fluorescence microscopy images as examples of criteria on metabolic activity assay of *Chroococcidiopsis* sp. UAM816 cells after 6 days of irradiation with UVR + PAR (A, B, C, D). A) DIC microscopy image where orange CTF crystals are visible in metabolically active cells (black arrows). B) eGFP filter set fluorescence image with revealing dead cells (GF+) (white arrows). C) Rhodamine filter set fluorescence image with revealing cells with phycobiliproteins and chlorophyll autofluorescence (white arrows) (PCHL+). D) HE rhodamine filter set fluorescence image of cells with phycobiliproteins and chlorophyll autofluorescence and weak CTF fluorescence (granulose red fluorescence) (blue dotted cells) (PCHL+/CTC+). E) HE rhodamine filter set fluorescence image of *Chroococcidiopsis* sp. UAM816 cells aggregate after only 3 days of irradiation with UVR + PAR. Cells with phycobiliproteins and chlorophyll red autofluorescence revealing still high metabolic activity (PCHL+/CTC+), yellow signal within the cyanobacteria cells. Scale bars = 10 μm.

In UAM813 strain (Fig. 6), 86.9-96.2 % of cells were active during the first 12 days of exposure to PAR. A final decrease in this active cells occurred after 15 days of exposure (90.1%). However, maximum of relative abundance of active cells under UVR+PAR light conditions was reached after 9 days of exposure (93.2%), exhibiting a progressive decrease after 12 and 15 days of exposure decaying below the starting point values (70.5%). In both PAR and UVR+PAR conditions, maximum values of damaged cells, 5.9% for PAR and 21.6% for UVR+PAR, were observed after 15 days of exposure, while the relative abundance of not viable cells reached its maximum after 6 days under PAR (4.7%) and after 15 days for UVR+PAR (7.9%).

**Figure 6.**
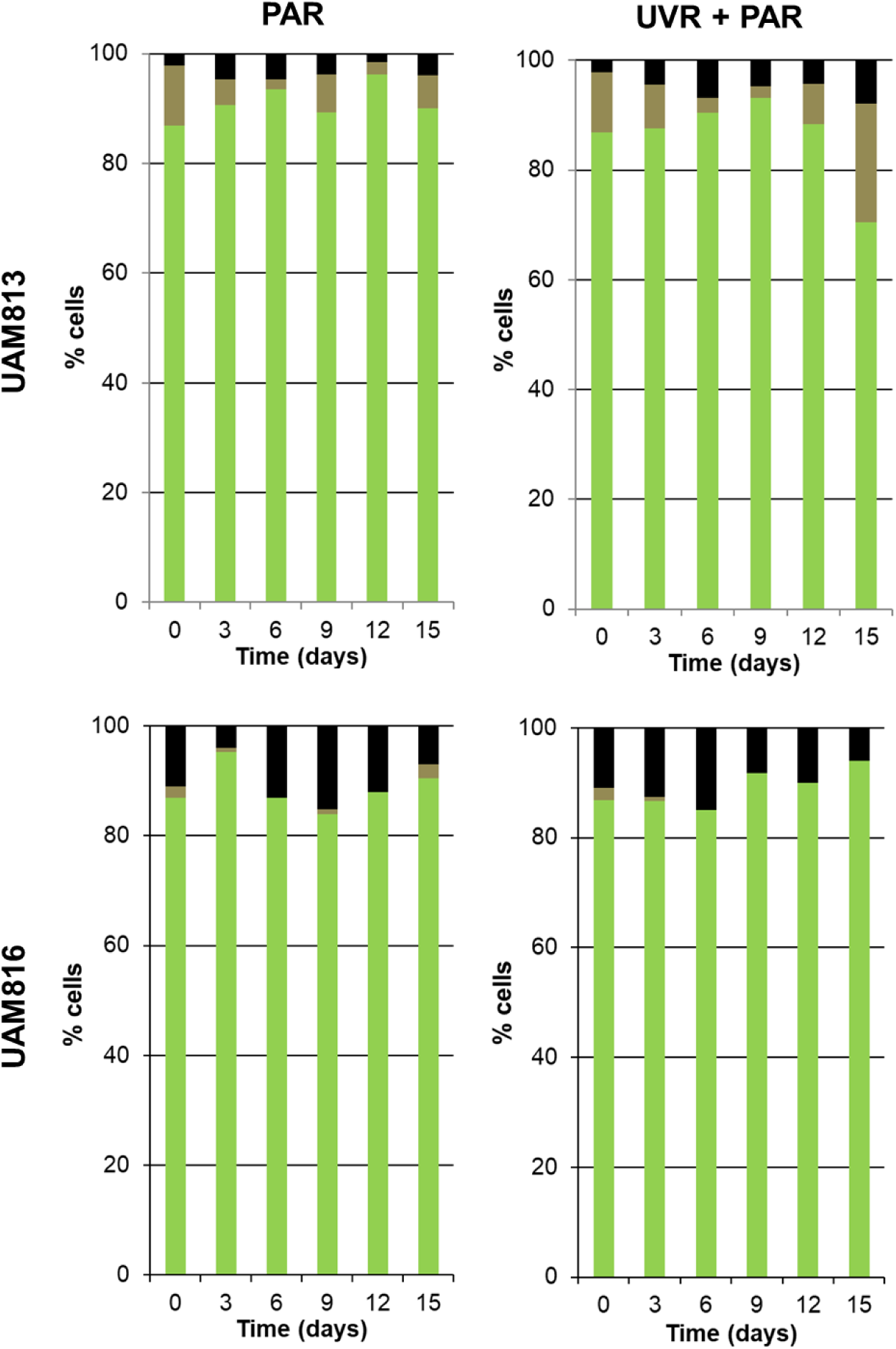
Metabolic activity of *Chroococcidiopsis* sp. UAM813 and UAM816 cells after irradiation with PAR (left graphs) or UVR + PAR (right graphs) for 15 days. Green: (CTC+/CHL+) active cells. Brown: (CHL+) damaged cells. Black: (GF+) dead cells.

The UAM816 strain cells metabolic activity (Fig. 6) exhibited a different behavior where the maximum values of active cells for PAR exposure were found after 3 days (95.2%) maintaining lower relative abundances during the following experimental times (86-90%). That maximum was reached after 15 days under UVR+PAR light conditions (94%) upon a progressive increase during the experiment. This progressive increase was accompanied by total absence of damaged cells under UVR+PAR, whereas under only PAR light for 15 days 2.5% of damaged cells were observed. The presence of death cells was detected during the whole experiment for both light conditions. When exposed to PAR for 9 days the culture experienced 15.1% (maximum ratio) of death cells. However, the quantity of death cells decreased to 7% after 15 days of experiment. A similar proportion was found when cells were exposed to UVR+PAR, reaching a maximum of 15% death cells after 9 days and decreasing to 6% at the end of the experiment.

#### Chroococcidiopsis *micromorphology and ultrastructure after its exposure to UVR and PAR*

Micromorphological and ultrastructural changes were examined in both UAM813 and UAM816 *Chroococcidiopsis* strains before UVA irradiation and after 9 days of exposure to UVR+PAR light (Figs. 7).

**Figure 7.**
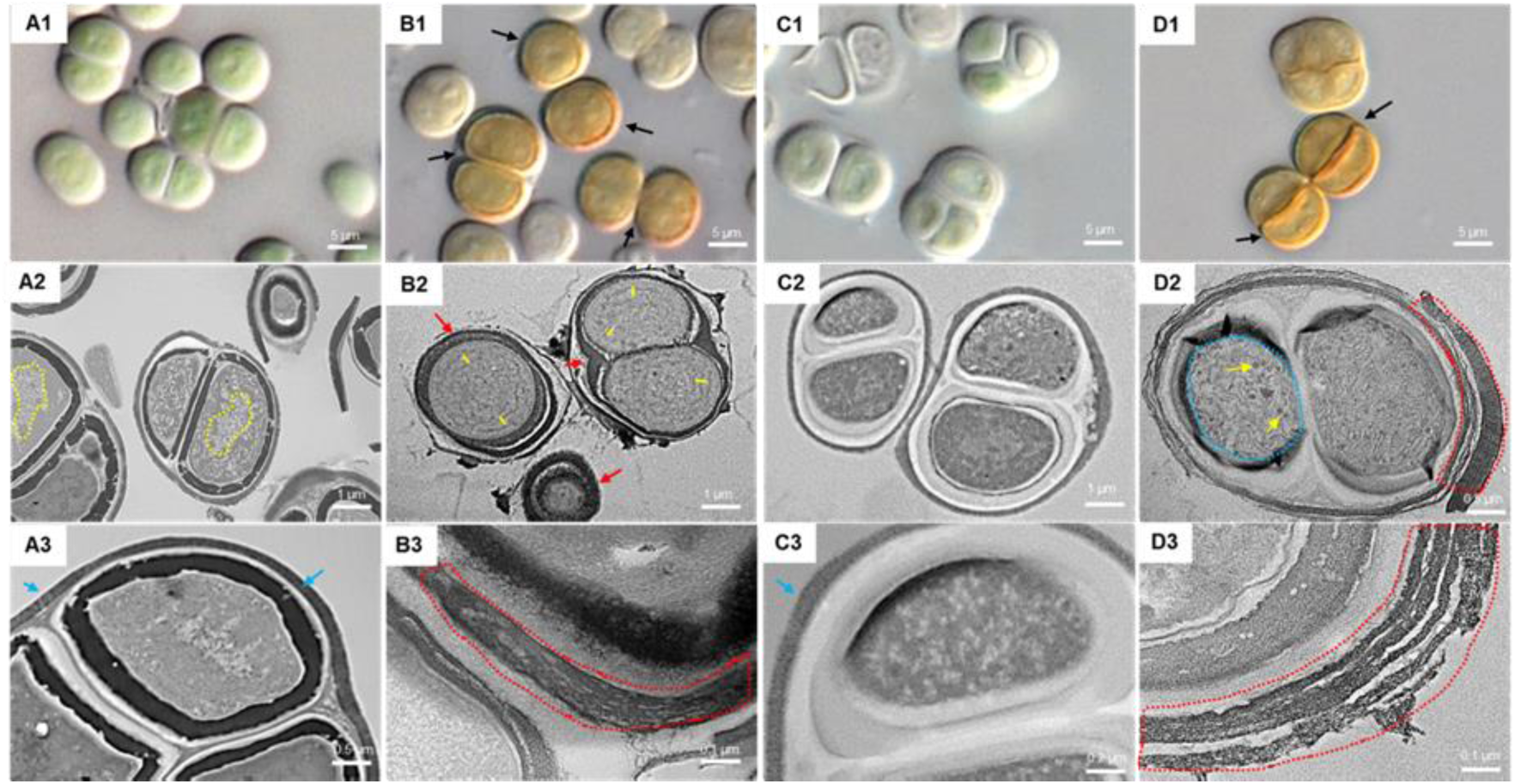
DIC microscopy and transmission electron microscopy images from *Chroococcidiopsis* sp. UAM813 (Series A and B) and UAM816 (Series C and D). **Series A**: Cells and aggregates at the beginning of the experiment. A1) Green *Chroococcidiopsis* sp. UAM813 cells. A2) TEM micrograph with cells exhibiting a visible nucleoid area (yellow dotted line). A3) TEM micrograph with an aggregate exhibiting a thin outermost fibrous layer (blue arrows). **Series B**: Cells and aggregates with maximum scytonemin content after 9 days of irradiation with UVR+PAR. B1) Brownish *Chroococcidiopsis* sp. UAM813 cells with higher scytonemin content on the edge of cells. B2) TEM micrograph with cells revealing a higher distance between thylakoids (yellow lines) and the presence of an electron dense outermost fibrous layer (red arrows). B3) TEM micrograph of the outer part of the cells from the same aggregate revealing a highly fibrous outermost layer (red dotted line). **Series C**: Cells and aggregates at the beginning of the experiment. C1) DIC image of green *Chroococcidiopsis* sp. UAM816 cells. C2) TEM micrograph of cells revealing the thylakoid arrangement within no irradiated cells. C3) TEM micrograph of *Chroococcidiopsis* sp. UAM816 cell revealing a slightly developed outermost fibrous layer (blue arrow). **Series D**: Cells and aggregates with maximum scytonemin content after 9 days of irradiation with UVR+PAR. D1) DIC image of brown *Chroococcidiopsis* sp. UAM816 cells with higher scytonemin content on the outer part of the cells. D2) TEM micrograph with cells revealing disaggregation of thylakoid membranes (blue dotted line), glycogen granules (dark spots pointed by yellow arrows) and a highly electron dense outermost fibrous layer (red dotted line). D3) TEM micrograph of the outer part of the cell revealing a highly fibrous outermost layer (red dotted line).

Changes of the cells color from light green to brownish were distinctly visible in UAM813 strain after UVR+PAR exposure (Figs. 7, A1-B1). Regarding the thylakoid placement in UAM813 strain cells after UVR+PAR exposure (Fig. 7, B2), an increase in the intra-thylakoid space was observed while the thylakoid membranes in cells not exposed to UVR (Fig. 7, A2) were positioned touching each other tightly showing a nucleoid area. A more developed electron dense outermost layer was observed in cells after their exposure to UVR+PAR (Fig. 7, B2) with a granulose and fibrous appearance (Fig. 7, B3) compared to the compact sheath observed in *Chroococcidiopsis* UAM813 cells that did not suffer UVR exposure (Fig. 7, A3).

UAM816 *Chroococcidiopsis* cells showed evident color differences too, when exposed to UVR+PAR for 9 days (Figs. 7, C1-D1). Intensive brownish was observed on the outer EPSs of the cell aggregates corresponding to an electron dense matrix within the EPSs (Fig. 7, B3-D3). Ultrastructural changes were found in different features as thylakoid arrangement showing the beginning of thylakoid membrane disintegration and glycogen granules along the thylakoids (Fig. 7, D2). The outermost fibrous layer observed in cells before the treatment (Fig. 7, C3) exhibited a more developed denser aspect with an assembly of various fibrous layers after its exposure to UVR+PAR for 9 days (Fig. 7, D3).

## 4. Discussion

This work provides a new insight into the behavior of cyanobacteria in endolithic communities under extreme solar radiation, as happens in the hyper-arid core of the Atacama Desert. Despite the sole development of lithobiontic microbial communities in endolithic habitats in different lithic substrates in this desert (39, 55–57) which act as a first line of defense against the damage provoked by high light exposure, the presence of second and third lines of cyanobacterial defense (39, 40, 42) points to the existence of specific, not previously characterized, adaptations to the harmful effects of high PAR and UVR, too.

In this work, two isolates from the extremotolerant genus of *Chroococcidiopsis*, widely distributed in the endolithic communities, were used to unravel the specific responses and adaptations to direct exposure to PAR and UVR. It was shown that both strains reveal specific differences matching their distinct microhabitat origin, cryptoendolithic from halite (UAM813) and chasmoendolithic from calcite (UAM816) substrates,

Both strains shared several common features, such as same genus – *Chroococcidiopsis*, same type of original habitat – endolithic -, and similar hyper-arid climatic conditions of their original habitat. Despite their similarities, evident differences were found in their response to direct light exposure that will be discussed below.

High response to PAR was observed in contrast to other studies when analyzing short-term (ROS accumulation) (6) and long-term response (scytonemin content) (46, 47, 58). In fact, both *Chroococcidiopsis* strains exhibited a lower short-term acclimation to PAR, compared to UVR+PAR light treatment. The same pattern was found regarding the long-term response of the UAM816 strain in both the scytonemin content and the metabolic activity tests. The high response to PAR light displayed in both cases could be explained by their original habitat, since by living in the endolithic microhabitat the direct and harmful exposure to solar radiation is avoided. This behavior was different in comparison to other studies where no PAR-induced oxidative stress was found in *Anabaena* (6) or *Nostoc* and *Fischerella* (58). The high response observed in the endolithic *Chroococcidiopsis* strains to PAR certainly support the requirement of a second and third line of defense against radiation, despite inhabiting endolithic microhabitats.

Concerning the short-term response, UAM813 strain exhibited a considerably lower acclimation to both light conditions where a subsequent increase of ROS during the 3 days of exposure with no signs of recovery was found. However, the UAM816 strain exhibited an acclimation to both light conditions, even better to UVR+PAR, with a similar response pattern to the one reported in *Anabaena* by He and Häder (6), although UAM816 acclimation started 24 hours earlier.

Long-term response to direct light should be explored considering three elements: scytonemin production, metabolic activity and ultrastructural changes. Both *Chroococcidiopsis* strains displayed clearly different responses in all three parameters. No severe ultrastructural damages were observed in the UAM813 strain when exposed to 9 days of direct light, although a visible increase of cover thickness was detected. This characteristic might be linked to the proportion of dead and damaged cells observed after 15 days of exposure, where both types of physiological status reached their maxima. This fact could explain the relatively low content of scytonemin in this strain during the experimental period. The UAM813 strain already showed its low capacity to cope with UVR in short-term exposure, and it seemed to happen again in long-term exposure. The relative scytonemin content reached its maximum after 9 days of exposure.

Subsequently, the low abundance of new cells able to produce scytonemin, as exhibited in the metabolic activity test, together with a slight increase in DW, due to the thickening of the cellular covers would maintain or slightly decrease the proportion between scytonemin and DW in the culture.

The long-term response of the UAM816 strain against UVR+PAR could be explained by its ultrastructural changes and metabolic activity. Higher ultrastructural damage could be observed after 9 days of exposure to UVR, coinciding with the experimental time where a major dead and damaged cell proportion was observed. The recovery of the physiological status after that point can be explained by the major capacity of the UAM816 strain to cope with this type of stress, as demonstrated in the short-term experiment expressed by ROS accumulation. Its capacity to recover and acclimate to the stressful conditions would allow an increase in the population leading to a subsequent decrease in the relative scytonemin content, since, thanks to its high acclimation capacity, growth would occur faster than the scytonemin production.

It was shown that acclimation capacity is strain-dependent, with significantly lower scytonemin content values in both *Chroococcidiopsis* strains from endolithic communities of the Atacama Desert than previously reported *Chroococcidiopsis* from desert crusts of the Vizcaíno Desert (Mexico) (46, 47).

The lower acclimation capacity of the UAM813 strain was observed in both short-term and long-term responses, which could be tightly linked to its original microhabitat. Since the *Chroococcidiopsis* UAM813 strain derived from a cryptoendolithic microhabitat, it would never be directly exposed to light, being always protected by the halite crust. This could explain its lower acclimation capacity to direct light exposure. On the contrary, the *Chroococcidiopsis* UAM816 strain comes from a chasmoendolithic microhabitat, thus being more exposed to direct PAR and UVR incidence by entrances of the fissures or cracks. The *Chroococcidiopsis* strains isolated from this microhabitat are therefore expected to be faster in the acclimation to light exposure.

There could be a linkage between each strain and its original microhabitat explained by a microhabitat specific environmental pressure. Thus, *Chroococcidiopsis* strains inhabiting certain endolithic microhabitats and lithic substrates could be absent from a different endolithic microhabitat and substrate in the same desert. This differential distribution could be explained by the possession of specific adaptations and the acclimation capacity of these organisms to the specific abiotic stresses occurring in the endolithic microhabitat they are inhabiting.

The third line of defense against intensive light displayed by these strains, such as the production of scytonemin, could have benefits, for both cyanobacteria and the entire endolithic community. The protection provided by scytonemin to cyanobacteria could be expected since these strains have shown extremely low growth rates in their original microhabitat (59) and each cell would suffer long exposure times that would lead to scytonemin production and accumulation. Furthermore, the accumulation of scytonemin could also provide protection against harmful radiation to the whole microbial community based on two considerations. On the one hand, the high stability of scytonemin (14, 40, 46, 47) deposited within EPS covers (42) as also observed in this study (Fig. 7). On the other, the desiccation for UAM813 strain, and salinity and desiccation for UAM816 strain conditions in the original endolithic microhabitat of these strains, known to promote the induction of scytonemin production (46, 47). Hence, the outcome of the combination of both conditions is a UV-screening effect over the whole endolithic microbial community that could enable its easier development through time, avoiding the harmful effects of extreme solar radiation.

## 5. Concluding Remarks

This is a pioneer study since it explores the response of cyanobacteria against UVR and PAR in cyanobacterial strains isolated from a place on Earth where one of the highest solar radiation levels have been detected, the hyper-arid core of the Atacama Desert.

The observed response of both *Chroococcidiopsis* strains to intensive light irradiance points at a strain specific selection on a microhabitat or substrate, related to the greater or lesser exposure to abiotic stresses. That selection would be based on the acclimation capacity and adaptation strategies displayed by different strains, confirming the statement “Everything is everywhere and the environment selects” (Baas-Becking, 1934). Everything meaning, the different *Chroococcidiopsis* strains, everywhere, the endolithic microhabitats of the hyper-arid core of the Atacama Desert, and selective environment, the slight differences in direct exposure, in this case to solar radiation, between lithic substrates and the type of endolithic microhabitat.

## Author contributions

MCC and JW designed, performed the research and conceived the original project. MCC, JW and AQ wrote the manuscript; MCC, JW and CA performed the microscopy; HMM performed HPLC analysis. All authors contributed to editing and revising the manuscript and approved this version for submission.

## Acknowledgments

This study was supported by grant PGC2018-094076-B-I00 from MCIU/AEI (Spain) and FEDER (UE). The work of MCC was supported by grant BES 2014-069106 from the Spanish Ministry of Science and Innovation (MCINN). The Instituto Estructura de la Materia-CSIC, Madrid, Spain and Dr. V. Souza-Egipsy is acknowledged for transmission electron microscopy services.

---

**Figure S1.**
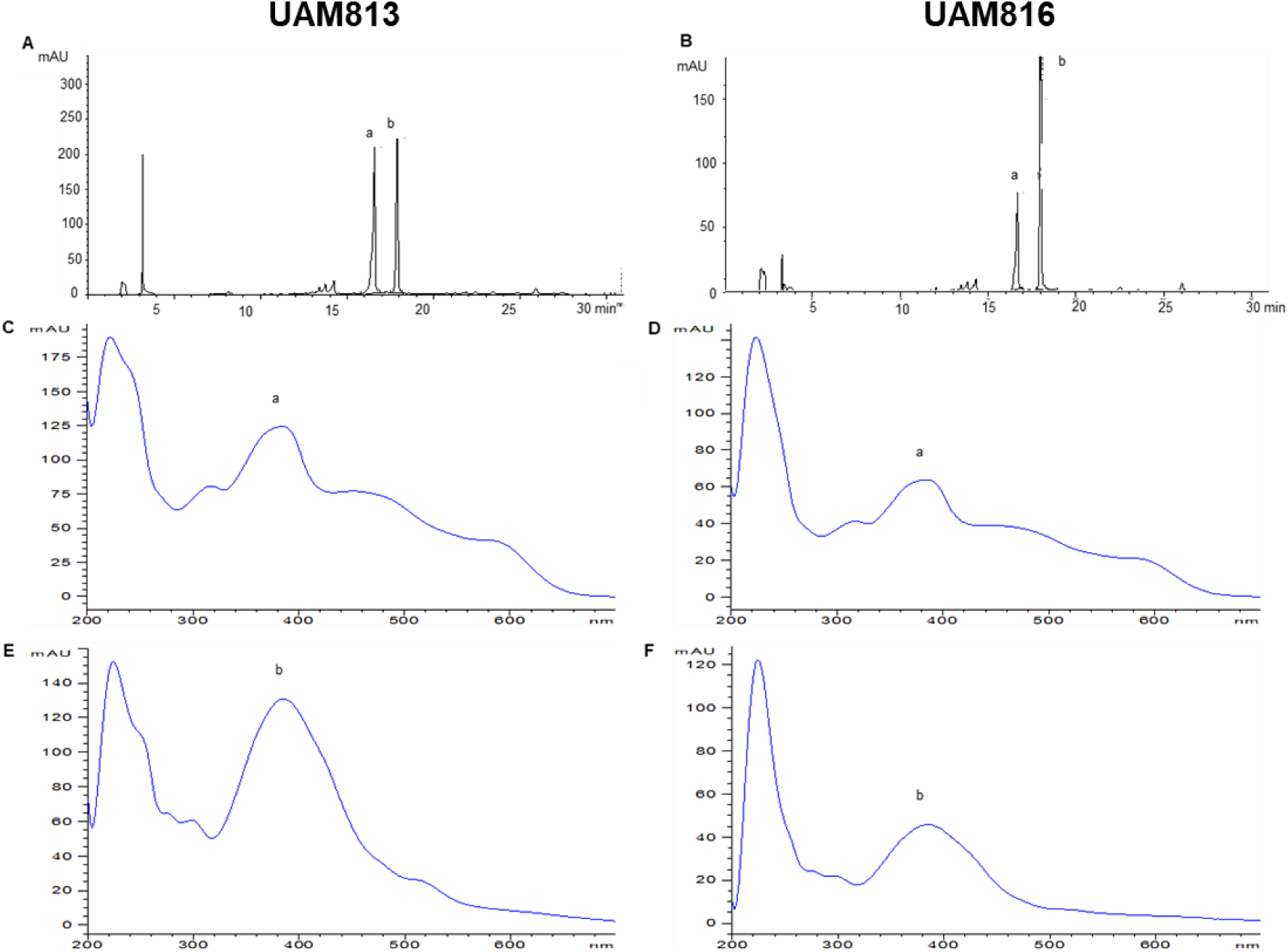
The HPLC chromatogram and absorption spectra of scytonemin extract of *Chroococcidiopsis* strains UAM813 (A, C, E) and UAM816 (B, D, F). A and B: The HPLC chromatogram of the reduced (a) and oxidized (b) scytonemin in UAM813 and UAM816. The absorption spectra of the reduced scytonemin of UAM813 (C) and UAM816 (D), and the oxidized scytonemin of UAM813 (E) and UAM816 (F)

